# The mouse *Klf1 Nan* variant impairs nuclear condensation and erythroid maturation

**DOI:** 10.1101/477059

**Authors:** Ileana Cantú, Harmen J.G. van de Werken, Nynke Gillemans, Ralph Stadhouders, Steven Heshusius, Alex Maas, Zeliha Ozgur, Wilfred F.J. van IJcken, Frank Grosveld, Marieke von Lindern, Sjaak Philipsen, Thamar B. van Dijk

## Abstract

Krüppel-like factor 1 (KLF1) is an essential transcription factor for erythroid development, as demonstrated by *Klf1* knockout mice which die around E14 due to severe anemia. In humans, >65 KLF1 variants, causing different erythroid phenotypes, have been described. The *Klf1 Nan* variant, a single amino acid substitution (p.E339D) in the DNA binding domain, causes hemolytic anemia and is dominant over wildtype KLF1. Here we describe the effects of the *Nan* variant during fetal development. We show that *Nan* embryos have defects in erythroid maturation. RNA-sequencing of the *Nan* fetal liver cells revealed that Exportin 7 (*Xpo7*) was among the ~780 deregulated genes. This nuclear exportin is implicated in terminal erythroid differentiation; in particular it is involved in nuclear condensation. Indeed, KLF1 *Nan* fetal liver cells had larger nuclei and reduced chromatin condensation. Knockdown of XPO7 in wildtype erythroid cells caused a similar phenotype. We conclude that reduced expression of XPO7 is partially responsible for the erythroid defects observed in *Nan* erythroid cells.

## INTRODUCTION

Erythropoiesis is the process of red blood cell production; defects in this process lead to anemia and thus insufficient oxygen delivery to tissues and subsequent organ failure. Therefore, the formation of red blood cells has to be tightly controlled during embryonic development and homeostasis in the adult.

KLF1 (previously known as EKLF) is a well-characterized, erythroid-specific transcription factor and one of the critical regulators of red blood cell maturation. KLF1 acts mainly as an activator and its target genes are involved in multiple processes of erythroid differentiation, including cell cycle regulation (1, 2), hemoglobin metabolism (3), and expression of membrane skeleton proteins (4, 5). The importance of KLF1 is illustrated by *Klf1* knockout embryos which die around E14 due to the lack of functional erythrocytes (6, 7). In contrast, heterozygous *Klf1*+/− mice survive into adulthood, showing that haploinsufficiency for KLF1 does not have a severe phenotype (8). KLF1 has a N-terminal transactivation domain and a C-terminal DNA binding domain, composed of three zinc fingers. They mediate specific DNA binding to 5’-CACCC-3’ motifs (9). Variants in human *KLF1* are found across the entire gene. The majority are missense variants in the three zinc fingers, which presumably alter the DNA binding/sequence recognition properties of KLF1. Mutations in KLF1 are associated with different phenotypes in humans (10), such as In(Lu) blood group (11), hereditary persistence of fetal hemoglobin (HPFH) (12), zinc protoporphyria (13), increased HbA2 (14), and congenital dyserythropoietic anemia (CDA) type IV (15).

The Neonatal anemia (*Nan*) mouse is an ethylnitrosourea (ENU)-induced semi-dominant hemolytic anemia model first described in 1983 by Mary Lyon (16), who positioned the variant on chromosome 8 (17). In 2010, *Klf1* was identified as the gene responsible for this phenotype, due to a single point mutation in the second zinc finger (p.E339D) (18, 19). While *Klf1 Nan* homozygous mice die around E10, KLF1 *Nan* heterozygous mice survive into adulthood displaying life-long hemolytic anemia. This indicates that the *Nan* variant affects the function of wildtype KLF1 protein, as this phenotype does not occur in *Klf1* haplo-insufficient mice (6-8, 18, 19). Indeed, the DNA binding properties of *Nan* KLF1 may be altered due to steric clash between the carboxyl group of p.339D and the methyl group of thymidine, resulting in the deregulation of a subset of target genes (19), although alternative models have been proposed (18).

Until now, research has focused on the effects of the *Nan* variant in adult mice (18-20). Given that KLF1 expression begins around E7.5 (21), it is of interest to investigate the impact of aberrant KLF1 activity during development. Here we investigated erythropoiesis during different stages of fetal development and observed impaired red blood cell maturation at E12.5, as assessed by flow cytometry analysis of the CD71 and Ter119 markers. Expression profiling of E12.5 fetal liver cells revealed 782 deregulated genes in *Nan* mutant samples including a host of known KLF1 targets such as Dematin and E2F2 (1, 4, 22). Intriguingly, the nuclear exportin XPO7, which has recently been implicated in nuclear condensation and enucleation during erythroid maturation (23), was one of the deregulated genes. XPO7 expression was significantly downregulated in the presence of the *Nan* variant erythroid progenitors, contributing to increased nuclear size. This partially explains the erythroid defects observed in *Nan* erythroid cells and provides a novel link between KLF1 and nuclear condensation.

## MATERIAL AND METHODS

### Mice

All animal studies were approved by the Erasmus MC Animal Ethics Committee. The mouse strains used were *Klf1 Nan* mutant (16) and *Klf1* knockout (6). Genotyping was performed by PCR using DNA isolated from toe biopsies. For *Nan* genotyping, the PCR product was digested with DpnII. Embryos were collected at E12.5, E13.5, E14.5 and E18.5; tail DNA was used for genotyping. Primer sequences are detailed in Supplementary Materials and Methods.

### Blood analysis

Peripheral blood was collected from the mandibular vein of adult mice, and standard blood parameters were measured with an automated hematologic analyzer (Scil Vet ABC, Viernheim, Germany).

### Cell culture and transduction

I/11 erythroid progenitors and primary mouse fetal liver cells were cultured as described (24). To induce differentiation of I/11 cells we used StemPRO-34 SFM (10639-011, life technologies) supplemented with 500 μg/mL iron-saturated transferrin (Scipac) and Epo (Janssen-Cilag, 10 U/mL). Lentiviral shRNAs targeting XPO7 were obtained from the Sigma MISSION shRNA library. The clones used are detailed in Supplementary Materials and Methods.

### RNA isolation and RT-qPCR analyses

RNA was extracted using TRI reagent (Sigma-Aldrich). To synthesize cDNA, 2 μg of RNA were used together with oligo dT (Invitrogen), RNase OUT (Invitrogen), and SuperScript reverse transciptase II (Invitrogen) in a total volume of 20 μL for 1 hour at 42°C. 0.2 μL of cDNA was used for amplification by RT-qPCR. Other experimental details and primer sequences are detailed in Supplementary Materials and Methods.

### Protein extraction and western blotting

Total protein extracts from mouse fetal liver cells were prepared according to (25). To visualize protein expression, cell lysates of ~3×10^6^ cells were loaded on 10% SDS-polyacrylamide gels for electrophoresis. The gels were transferred to nitrocellulose blotting membrane 0.45 μm (10600002, GE Healthcare) and probed with specific antibodies. Membranes were stained for Tubulin (T5168, Sigma-Aldrich) as loading control, and for XPO7 (sc390025, Santa Cruz).

### Flow cytometry, cell sorting, enucleation- and cell morphology analysis

These procedures are described in detail in Supplementary Materials and Methods.

### RNA-sequencing and analysis

RNA-seq was performed according to manufacturer’s instructions (Illumina; San Diego, CA, USA), as described(26). The sequenced reads were mapped against the mouse genome build mm10 using TopHat 2.0.6 (27) with the transcriptome gene annotation of Ensembl v73 (28). Further details of the bioinformatics analyses are described in Supplementary Materials and Methods.

### Chromosome Conformation Capture Combined with high-throughput Sequencing (4C-seq) and data analysis

4C-seq experiments were carried out as described (29, 30). Briefly, 4C-seq template was prepared from E13.5 fetal liver or fetal brain cells. In total, between 1 and 8 million cells were used for analysis. Further experimental details and of the bioinformatics analyses are described in Supplementary Materials and Methods.

### Statistical tests

Statistical analysis of blood parameters was performed by using analysis of variance with Bonferroni correction; flow cytometry data and gene expression results were analyzed by using Mann-Whitney tests. Excel 2010 (Microsoft, Redmond, WA) was used to draw the graphs. Values plus or minus standard deviation are displayed in all figures.

## RESULTS

### Characterization of *Nan* fetal liver cells

The effect of the KLF1 *Nan* variant has been studied in adult mice (18-20), but data on its effect during gestation is limited. Hence, to study this variant during embryonic development, we used a *Nan* mouse model carrying one mutant allele (*Nan*/+, from now on called *Nan*). At E12.5, E14.5, and E18.5, *Nan* embryos were paler than wildtype littermates, indicating anemia, but otherwise looked phenotypically normal. Flow cytometry analysis of E12.5, E14.5, and E18.5 fetal liver cells used the Kit, CD71, Ter119 and CD44 markers to trace red blood cell differentiation. A severe downregulation in expression of the Ter119 marker was observed at all three stages (Figure 1A,B). The CD71/Ter119 double-positive population was significantly decreased in the *Nan* samples, while the CD71 single-positive population showed an increase. No significant differences were observed for Kit and CD44 in the *Nan* variant (Supplementary Figure 1A,B). In addition, similar results were obtained when assaying embryonic blood, with Ter119 being highly downregulated (Figure 1C,D). These results indicate that *Nan* embryos display delayed erythroid maturation compared to wildtype controls. This is in line with the observation that a higher percentage of cells is positive for CD71 in adult blood (Supplementary Figure 2A,B), indicative of higher percentage of circulating reticulocytes (19). Consistent with this notion, analysis of standard blood parameters revealed a significant increase in red cell distribution width (RDW) in the *Nan* mice (Supplementary Figure 2C). Furthermore, we observed minor, yet significant, decreases in RBC (red blood cell), HGB (total hemoglobin), HCT (hematocrit), MCH (Mean Corpuscular Hemoglobin), MCHC (Mean Corpuscular Hemoglobin Concentration) values. Interestingly, when comparing *Nan* E14.5 fetal liver cytospins to wildtype controls, we observed a marked increase in the average size of the erythroid cells and their nuclei (Figure 1E). Taken together, these data show that erythroid maturation is impaired in *Nan* animals.

**Figure 1.**
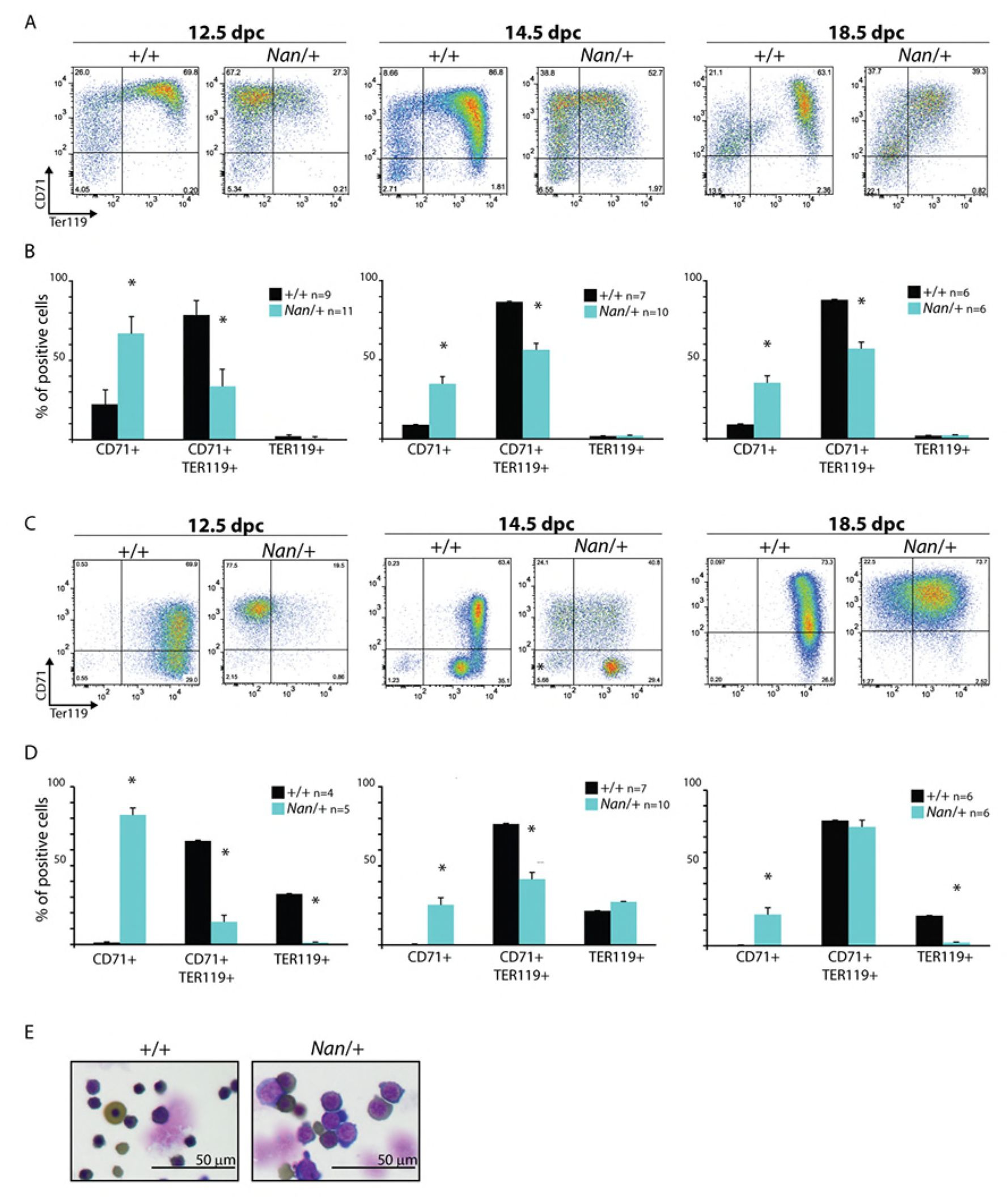
Flow cytometry analysis of erythroid cells isolated from *Nan* embryos. (A) Examples of flow cytometry profiles of CD71 and Ter119 staining of E12.5, E 14.5 and E18.5 wildtype and *Nan* mouse fetal livers. (B) Quantification of CD71+, CD71+ Ter119+ and Ter119+ populations. n indicates the number of embryos. * indicates *p value* <0.01. (C) Examples of flow cytometry profiles of CD71 and Ter119 staining of E12.5, E 14.5 and E18.5 wildtype and *Nan* mouse fetal blood. (D) Quantification of CD71+, CD71+ Ter119+ and Ter119+ populations. n indicates the number of embryos. * indicates *p value* <0.01. (E) Cytospins of E14.5 wildtype and *Nan* mouse fetal liver cells stained with May Grünwald-Giemsa and O-dianisidine.

**Figure 2.**
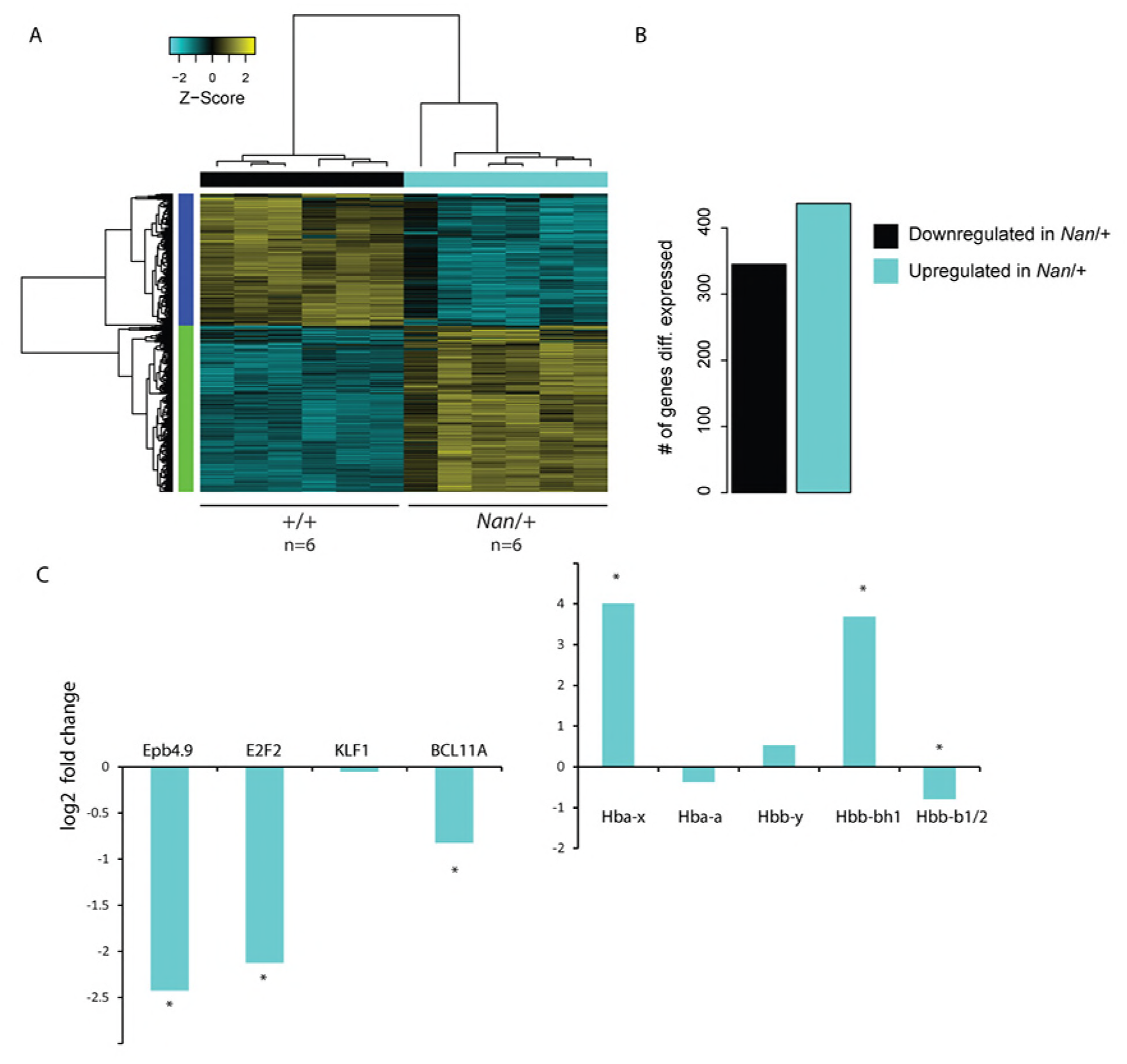
RNA-seq analysis of wildtype and *Nan* fetal liver cells. (A) Hierarchical clustered heat map with scaled Z-score color key of normalized counts of 782 differentially expressed genes in 6 WT (+/+) and 6 *Nan* (*Nan*/+) E12.5 fetal liver samples. Samples with the same genotype are indicated by black (WT) and cyan (*Nan*) horizontal bars; gene clusters are indicated by green (upregulated in *Nan*) and purple (downregulated in *Nan*) vertical bars. False discovery rate [FDR] <0.01, fold-change equal or greater than 1.5. (B) Schematic representation of the number of downregulated and upregulated genes in the *Nan* E12.5 fetal livers. (C) Log2 values of fold-change for selected genes. * indicates FDR <0.01.

### Identification of deregulated genes in E12.5 *Nan* fetal livers

In order to identify genes that are affected by the *Nan* variant, a genome-wide RNA-seq was performed on samples derived from E12.5 *Nan* and wildtype fetal livers (*N*=6 each), as at this stage the fetal liver is mainly composed of erythroid cells. 782 genes appeared to be deregulated in the *Nan* mutants (false discovery rate [FDR] <0.01, absolute fold change equal or greater than 1.5), of which 437 were upregulated and 345 downregulated (Figure 2A,B and Supplementary Table 1). Strikingly, even though KLF1 has been mainly described as a transcriptional activator, the majority of the deregulated genes displayed increased activation in the *Nan* erythroid cells. We postulate that this might be due to secondary effects of KLF1 on other transcriptional regulators and/or aberrant activity of KLF1 *Nan*. To validate the data, we checked the expression of *Epb4.9* and *E2f2*, genes known to be down-regulated in *Nan* erythroid cells (19) (Figure 2D, left panel and Supplementary Figure 3). Indeed, a significant decreased expression of the transcripts of these two genes was detected in *Nan* fetal livers. Moreover a significant 2-fold down-regulation of BCL11A, a known target of KLF1 (12, 31), was observed indicating that the KLF1 *Nan* variant affects its expression (Figure 2D, left panel and Supplementary Figure 3). Given the role of BCL11A and KLF1 in globin switching, the expression levels of the β-like globin genes were checked; the embryonic *Hbb-bh1* gene was upregulated and the KLF1 target gene *Hbb-b1* was downregulated, consistent with previous reports (19). In addition, the embryonic *Hba-x* gene was upregulated in E12.5 *Nan* fetal livers (Figure 2D, right panel and Supplementary Figure 3). Collectively, these data are in accordance with the notion that intact KLF1 fulfils a crucial role in developmental regulation of globin gene expression (8) and deregulation of embryonic globin expression in adult *Nan* mice (19).

**Figure 3.**
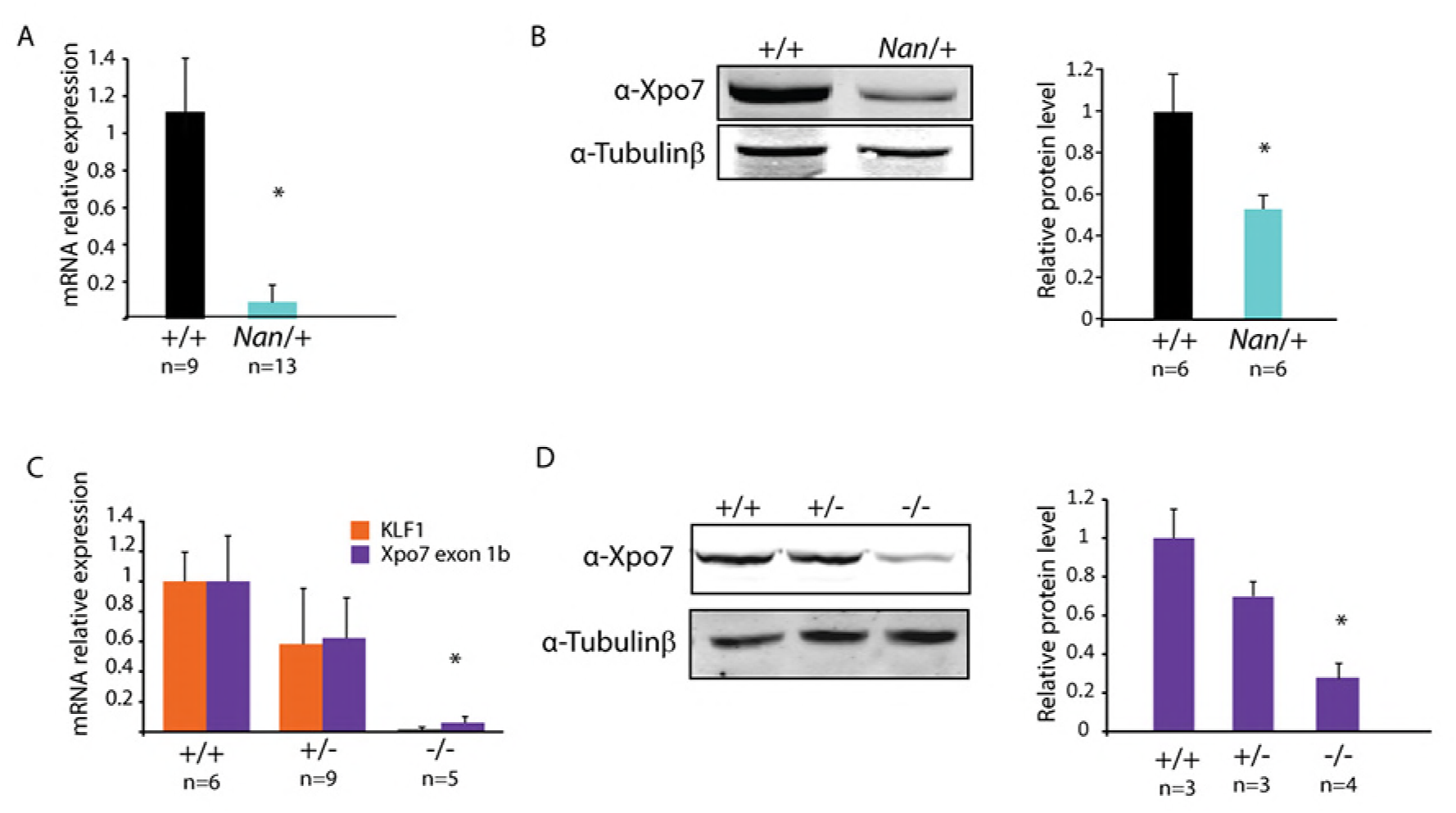
XPO7 expression in wildtype, *Nan* and *Klf1* knockout fetal liver cells. (A) XPO7 mRNA relative values in wildtype and *Nan* E14.5 fetal livers. * indicates *p value* <0.01. n indicates the number of embryos. (B) Western blot analysis of XPO7 protein in wildtype and *Nan* E14.5 fetal livers and quantification. β-tubulin was used as loading control. * indicates *p value* <0.01. n indicates the number of embryos. (C) KLF1 and XPO7 mRNA relative expression values in wildtype, KLF1 heterozygotes and KLF1 knockout E13.5 fetal livers. * indicates *p value* <0.01. n indicates the number of embryos. (D) Western blot analysis of XPO7 protein in wildtype, KLF1 heterozygotes and KLF1 knockout E13.5 fetal livers and quantification. β-tubulin was used as loading control. * indicates *p value* <0.01. n indicates the number of embryos.

### The nuclear exportin XPO7 is downregulated in *Nan* erythroid cells

The expression of the *Xpo7* gene, encoding a nuclear exportin, was prominently downregulated in *Nan* E12.5 fetal livers (~4-fold decrease; Figure 2A). This raised our interest since XPO7 was recently implicated in terminal erythroid differentiation, as a protein involved in enucleation (23). To corroborate the RNA-seq data, XPO7 expression in *Nan* E14.5 fetal livers was reduced by approximately 90% in the *Nan* samples as determined by RT-qPCR (Figure 3A). Transcripts using the canonical first exon were barely detectable in all samples (not shown). Importantly, XPO7 protein levels were also reduced in *Nan* fetal liver cells (Figure 3B). To investigate whether XPO7 expression is dependent on KLF1, XPO7 mRNA and protein levels were measured in *Klf1 null* erythroid cells. In E13.5 *Klf1 null* fetal livers (6) expression of XPO7 mRNA and protein is significantly reduced (Figure 3C,D). Remarkably, downregulation of XPO7 was also observed in *Klf1* wt/ko fetal livers, although to a lesser extent than observed in *Klf1 null* fetal livers (Figure 3C,3D). Thus, similar to BCL11A (12, 31), activation of XPO7 by KLF1 is dose-dependent. In agreement with the notion that KLF1 is a direct activator of the *Xpo7* gene, KLF1 binds to the canonical promoter and first intron of the *Xpo7* gene in mouse (32) and human (33) erythroid cells (Supplementary Figure 4).

**Figure 4.**
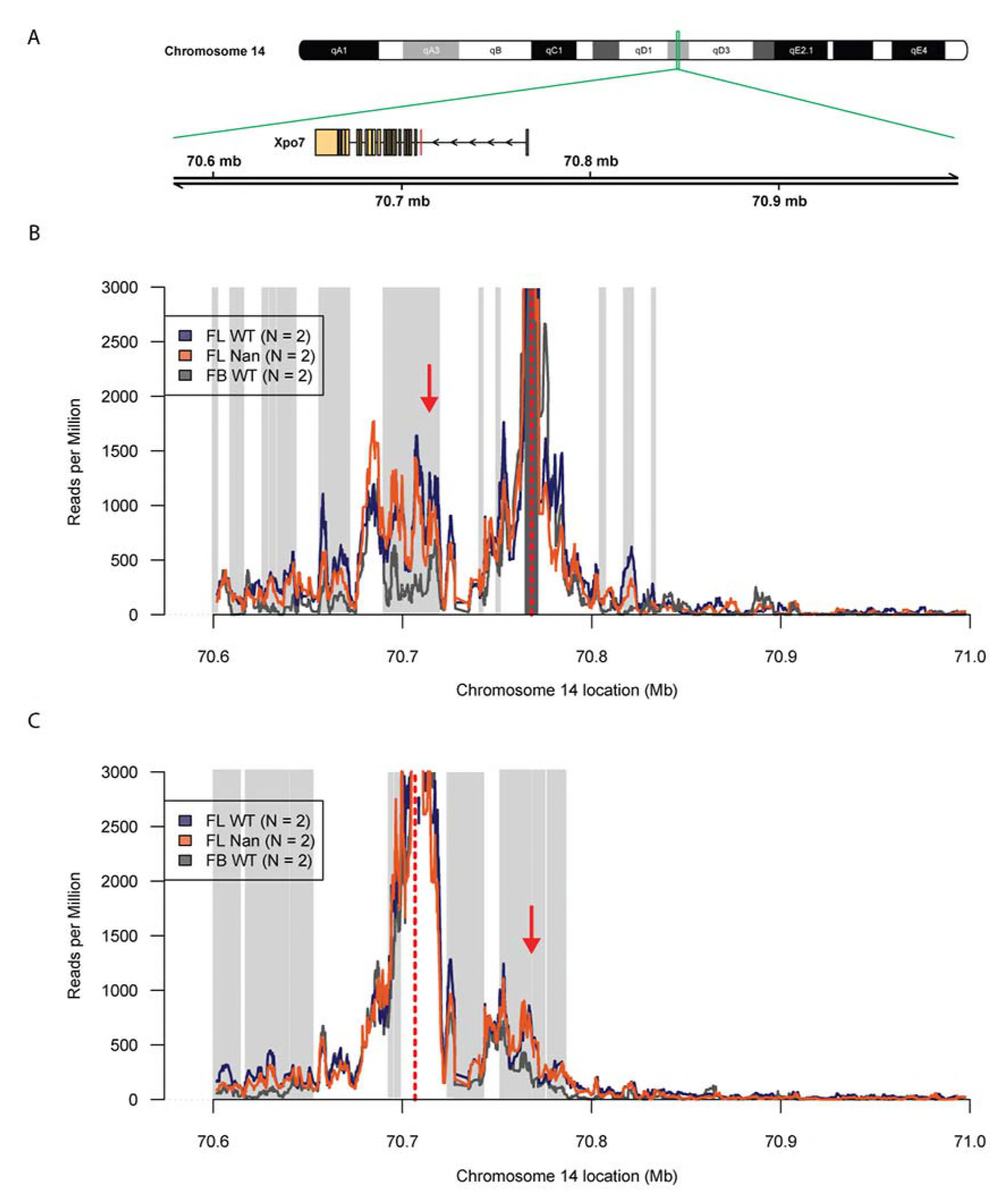
4C-seq analysis of the *Xpo7* locus. (A) Schematic representation of chromosome number 14. The green box indicates the zone where the *Xpo7* gene resides. The RefSeq mm10 Xpo7 gene is indicated by rectangles (exons) and arrows (introns) that point to the direction of transcription. The location is indicated in Mega basepairs (Mb). The erythroid-specific first exon is indicated by a red box. (B-C) 4C-seq representation of the chromosome contact frequencies detected using the canonical promoter of *Xpo7* (B) and the region of the erythroid specific *Xpo7* exon (C) as viewpoints. The mean of a running windows of 21 restriction fragment-ends of the median value of the biological replicates with a maximum of 3000 are indicated by colored lines. Loci with a statistically significant (FDR <0.05) higher contact frequencies and reads per million >250 in wildtype fetal liver compared to the fetal brain are indicated by light grey boxes. Loci with a statistically significant (FDR <0.05) higher contact frequencies and reads per million >250 in fetal brain compared to the wildtype fetal liver brain are indicated by dark grey boxes. The red dotted line indicates the view point and the red arrow the position of the loop. Purple, KLF1 +/+ fetal liver; Orange, KLF1 *Nan/*+ fetal liver; Grey, *KLF1* +/+ fetal brain.

### The chromatin conformation of the *Xpo7* locus is not affected in *Nan* erythroid cells

Since KLF1 is required to form an active chromatin hub in the β-globin locus (34), 4C-seq experiments were performed on the *Xpo7* locus in E13.5 wildtype fetal livers and fetal brains and *Nan* mouse fetal livers (Figure 4A). The canonical promoter of *Xpo7* was used as viewpoint to investigate potential changes in chromatin conformation (Figure 4B). Interestingly, a loop was identified between the canonical promoter of *Xpo7* (situated at the beginning of exon 1a) and the exon that produces the erythroid-specific form of Xpo7 (exon 1b), indicating that these two regions are in spatial proximity in erythroid cells (Figure 4B), whereas this loop has lower contact frequencies in fetal brain. However, few local changes in the chromatin conformation were found between wildtype and *Nan* samples (Figure 4B). The experiment was repeated using the erythroid-specific promoter as view point. This confirmed the results obtained with the canonical promoter as viewpoint. (Figure 4C). We suggest that this loop might recruit transcription factors binding to the area of the canonical promoter to the vicinity of the erythroid promoter, thereby facilitating expression of the erythroid-specific *Xpo7* transcript.

### *Nan* mouse fetal liver cells present defects in nuclear condensation

Since Xpo7 has been implicated in enucleation of erythroid cells *in vitro* (23) enucleation in *Nan* fetal livers was analyzed. This was quantified in E14.5 fetal livers by flow cytometry using the erythroid marker Ter119 and Hoechst-33342 as a nuclear stain. Similar percentages of enucleated cells were observed between *Nan* and control fetal liver samples (Figures 5A,B). Similar results were obtained with E12.5 and E18.5 fetal liver cells (data not shown). To check whether the flow cytometry analysis could indeed discriminate nucleated from enucleated cells, we sorted the Hoechst-positive and Hoechst-negative populations and prepared cytospins. This showed all the Hoechst-negative cells identified by flow cytometry to indeed be enucleated (Figure 5C). In addition, assessing enucleation levels of mouse fetal liver cells cultured in proliferative medium and in differentiation medium from embryos at E12.5 and E14.5 by flow cytometry, similar ratios of nucleated *versus* enucleated cells were found in control and *Nan* samples (Supplementary Figure 5A,B). Nevertheless, a striking increase in the percentage of large cells in the fetal livers of the *Nan* embryos was observed. Quantification of cell size using flow cytometry revealed a significant increase in average cell size at E12.5, E14.5 and E18.5 in the *Nan* samples (Figure 5D). In line with this finding the nuclear area of the *Nan* fetal liver cells was significantly increased when compared to control fetal liver cells in cytospin slides stained with the nuclear dye Hoechst 33342 (Figure 5E). These data are consistent with the notion that XPO7 is involved in nuclear condensation, a process that precedes enucleation. However, despite the impaired nuclear condensation, the cells still are still able to undergo enucleation as we observed similar ratios of nucleated *versus* enucleated cells in control and *Nan* fetal liver cells.

**Figure 5.**
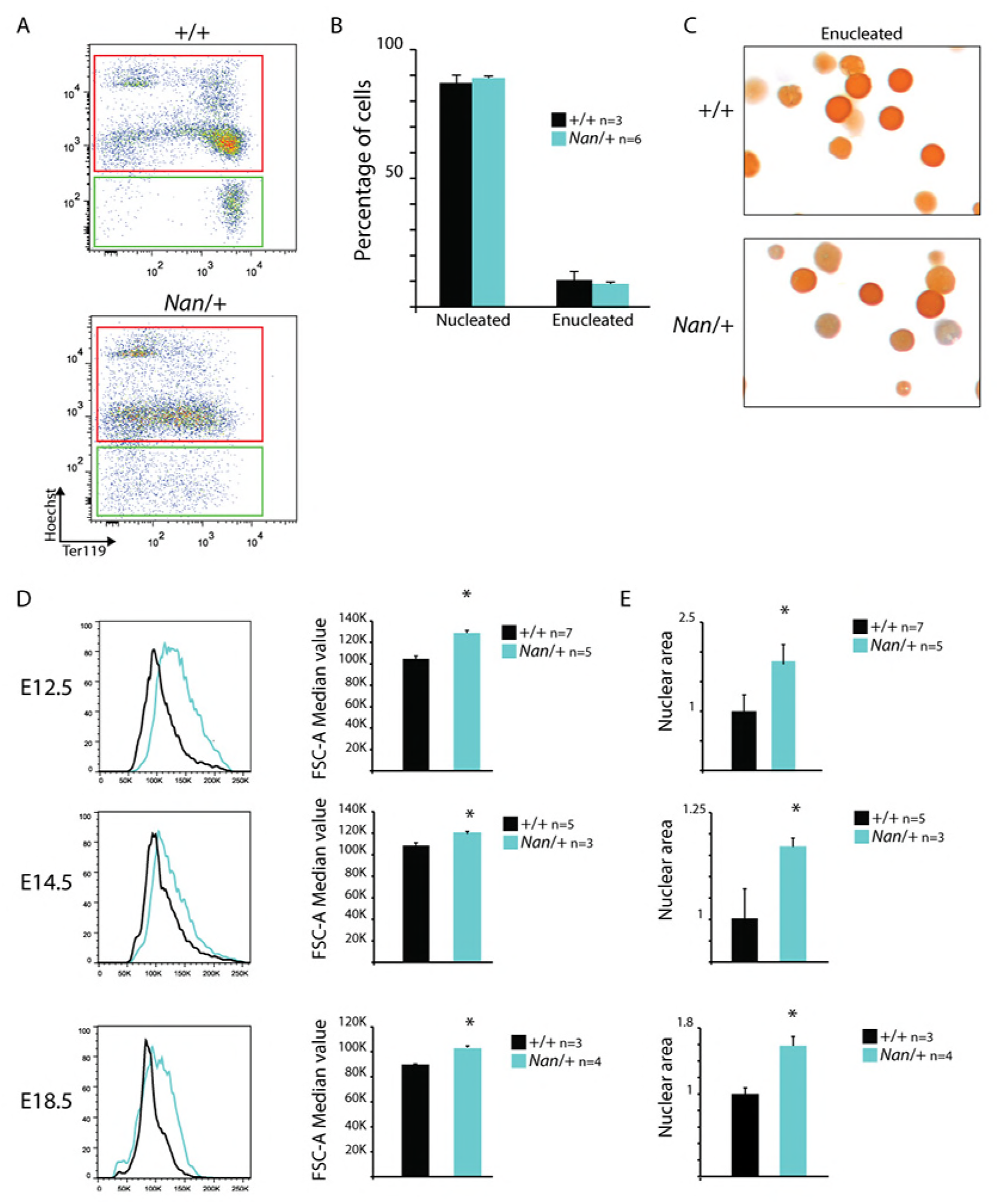
Analysis of enucleation and cell size of *Nan* fetal liver cells. (A) Gating strategies of Hoechst- and Ter119-stained E14.5 fetal liver cells. Red, Hoechst+ population; Green, Hoechst-population. (B) Quantification of the number of nucleated (Hoechst+) and enucleated (Hoechst-) cells. n indicates the number of embryos. (C) Cytospins stained with May Grünwald-Giemsa and O-dianisidine of Hoechst-wildtype and *Nan* sorted populations. (D) Representative FSC-A value flow cytometry plots of E12.5, E14.5 and E18.5 wildtype and *Nan* fetal liver cells and quantification. * indicates *p value* <0.01. n indicates the number of embryos. (E) Relative nuclear area size quantification of E12.5, E14.5 and E18.5 wildtype and *Nan* fetal liver cells. * indicates a p value <0.01. n indicates the number of embryos.

### XPO7 knock down in I/11 cells mimics the phenotype of *Nan* cells

The role of XPO7 in erythroid differentiation was further analyzed by knocking down XPO7 in the factor-dependent immortalized mouse erythroid cell line I/11 (35). Using three different shRNAs, an efficiency of ~70% knockdown was reached as shown by Western blot (Figure 6A). Before differentiation a minor difference in expression of the surface markers CD71 and Ter119 and no difference in cell size between the control and the knockdown cells was observed (data not shown). In contrast, upon transfer to differentiation medium, the maturation of XPO7 knockdown cells was impaired, as shown by CD71 and Ter119 flow cytometry analysis (Figure 6B), and the average cell size was increased (Figure 6C). In addition, using an ImageStream flow cytometer showed the mean and median size of the nuclear area to be increased upon XPO7 knockdown when the cells were cultured under differentiation conditions. This is consistent with the notion that XPO7 is required for nuclear condensation during terminal erythroid differentiation. Collectively, these findings indicate that XPO7 is partially responsible for the phenotype of *Nan* mice, establish that the *Xpo7* gene is a novel erythroid target gene of KLF1, and that nuclear condensation is a process previously unrecognized to be regulated by KLF1.

**Figure 6.**
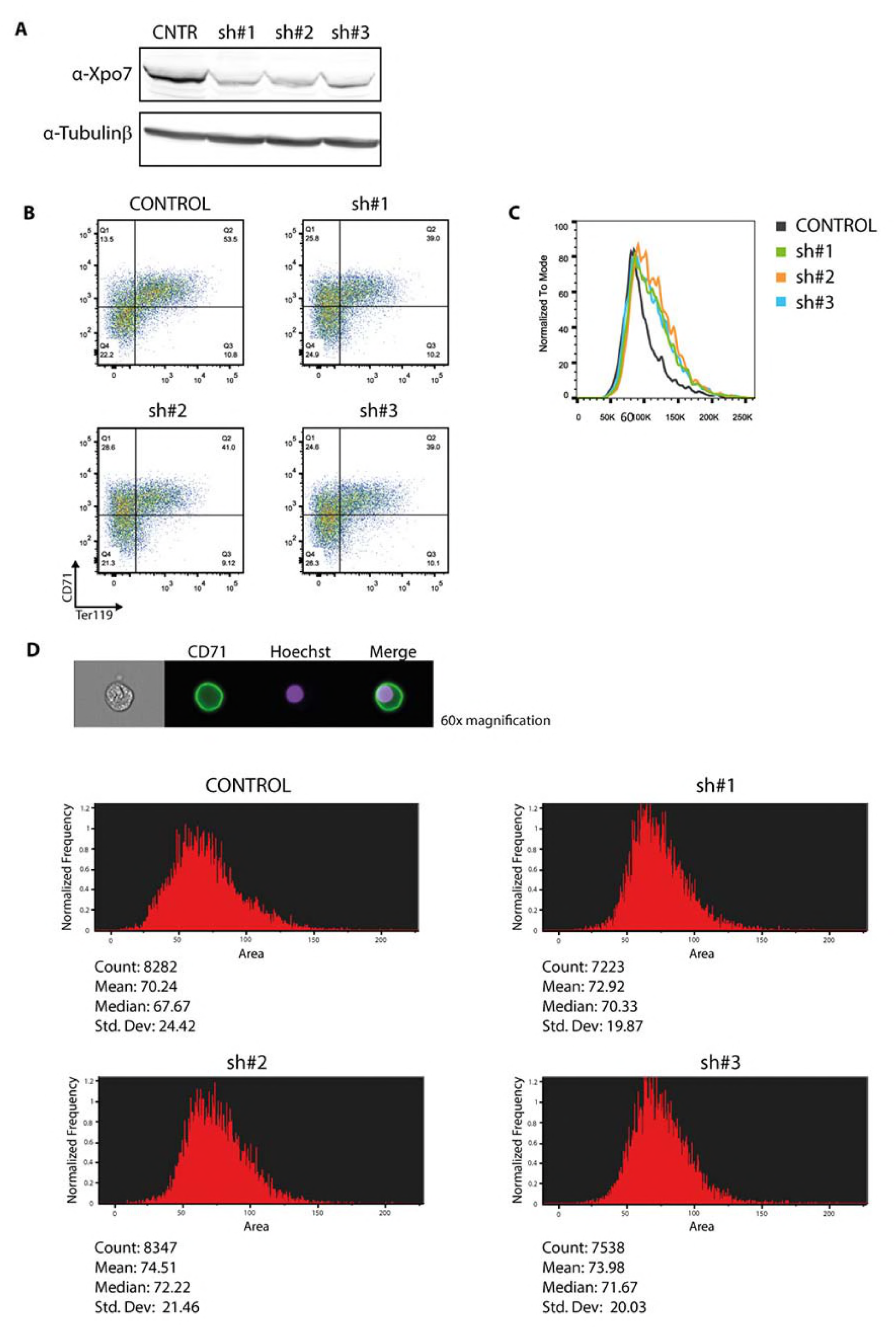
XPO7 knockdown in I/11 immortalized mouse erythroid progenitor cells. (A) Western blot analysis indicating the efficiency of XPO7 knockdown using 3 different shRNAs. β-tubulin was used as loading control. (B) Example of flow cytometry profiles of CD71 and Ter119 staining of I/11 cells transduced with either control, sh#1, sh#2 or sh#3 lentiviruses in differentiation conditions. The percentage of cells in the CD71/Ter119 double-positive quadrant is 50.2±1.92 for control cells and 43.5±3.0 for XPO7 knockdown cells (p=0.039, four independent experiments). (C) Representative FSC-A value flow cytometry plots of I/11 cells transduced with either control, sh#1, sh#2 or sh#3 lentiviruses in differentiation conditions. (D) ImageStream area quantification (arbitrary units) of I/11 cells transduced with either control, sh#1, sh#2 or sh#3 lentiviruses in differentiation conditions. The total number of cells counted, the mean, the median and the standard deviation are shown below the histograms. On top a representative cell from the control sample is depicted.

## DISCUSSION

Erythropoiesis is a complex process that involves many players whose coordinated activity ensures the production of functional red blood cells. One of these players is KLF1, a transcription factor with multiple roles during terminal erythroid differentiation. Firstly, it is essential for globin regulation, in particular for direct activation of β-globin (6, 7). In addition, it acts as a master regulator of genes activated during differentiation of red blood cells, such as membrane proteins, heme synthesis enzymes and cell cycle regulators (2, 4, 22). Hence, it comes as no surprise that *Klf1* knockout embryos die due to severe anemia, and that the phenotype is not rescued by exogenous expression of a β-like globin gene (36). Accordingly, *KLF1* variants can lead to diverse phenotypes in humans (10). One example is a missense variant in the second zinc finger of human KLF1 (p.E325K) that causes CDA type IV (15). This variant is believed to affect binding of KLF1 to its target genes thereby exerting a dominant-negative effect on wildtype KLF1 protein. Similar effects have been described for the *Nan* mouse model. These mice have a missense variant, p.E339D, in a position homologous to that of the human CDA type IV variant (18, 19). Studies on the effect of the *Nan* variant in adult mice have revealed that these animals display life-long anemia (18-20).

In this paper we present our findings on the effects of the *Nan* variant on definitive fetal erythropoiesis and show that erythroid maturation is impaired in *Nan* fetal livers at E12.5, E14.5 and E18.5. We identified 782 differentially expressed genes in *Nan versus* control E12.5 fetal livers. In agreement with a previous report on erythropoiesis in adult *Nan* mice (19), the expression of globin genes is altered in *Nan* fetal livers. In particular, the upregulation of embryonic βh1 globin can be explained by the significantly lower expression of BCL11A in *Nan* embryos, which normally suppresses βh1 expression (37). *Xpo7*, encoding a nuclear exportin, was one of the most significantly downregulated genes. It caught our attention since a recent paper described that *Xpo7* is required for nuclear condensation and enucleation during terminal erythroid differentiation *in vitro* (23). In addition, the observation that XPO7 expression was also reduced in *Klf1* knockout fetal livers indicated that the *Xpo7* gene might be a direct target of KLF1. Supporting this notion, data mining of ChIP-seq results revealed that KLF1 binds to the *Xpo7* locus in mouse (32) and human (33) erythroid cells. Collectively, these data suggested that, similar to the β-globin locus (34), KLF1 might have a role in the spatial organization of the *Xpo7* locus. 4C-seq analysis of the *Xpo7* locus demonstrated that it adopts a different conformation in fetal liver cells compared to fetal brain cells. The presence of the *Nan* variant doesn’t appear to mediate any major changes in the chromatin conformation of the *Xpo7* locus. We note that the promoter of the *Xpo7* gene contains so-called ‘category II’ KLF1 binding sites(19) which are recognized by wildtype KLF1 only. The presence of such ‘category II’ sites is a hallmark of downregulated genes in *Nan* erythroid cells. This suggests that in *Nan* cells wildtype KLF1 is still able to bind to the *Xpo7* promoter and organize the erythroid-specific 3D conformation of the locus, but is unable to activate transcription efficiently. An interesting observation is the presence of a loop between the promoter of the canonical *Xpo7* promoter (in front of exon 1a) and the erythroid-specific promoter (in front of exon 1b), which is absent in the fetal brain control. This loop is likely the consequence of recruitment of the two promoters to the same transcription factory (38). Previous work has shown that XPO7 knockdown in cultured mouse fetal liver cells impairs chromatin condensation and enucleation during terminal erythroid differentiation (23). Although in *Nan* mice enucleation still occurs, the reduced XPO7 expression due to the *Nan* variant impairs chromatin condensation during terminal erythroid differentiation. We propose that this contributes to the maturation defects of *Nan* erythrocytes in fetal and adult definitive erythropoiesis. Indeed, knockdown of XPO7 in immortalized mouse erythroblasts cells leads to impaired maturation of the cells, evident by dysregulation of the flow cytometry markers CD71 and Ter119 and the presence of larger cells with larger nuclei in the cultures. Our data are in reasonable agreement with the recent publication on the role of XPO7 in erythroid maturation (23). It is important to keep in mind that we cannot compare the levels of XPO7 protein between our system and that of Hattangadi et al. (23). An emerging question is how *Nan* cells manage to enucleate in the presence of reduced levels of XPO7. One possibility is that the level of XPO7 present *in vivo* in *Nan* mice might suffice for correct enucleation of the erythroblasts but still affects nuclear condensation. Alternatively, downregulation of XPO7 might just slow down nuclear condensation, but the cells eventually manage to shed their nucleus when condensation is completed. Lastly, a protein with a role similar to that of XPO7 may substitute for it, thus enabling enucleation. We favour a scenario in which chromatin condensation is crucial for enucleation (39-41), with XPO7 as an important effector. It is not clear whether enucleation can happen before nuclear condensation is completed. Our study suggests that impaired nuclear condensation contributes to the erythroid maturation defects observed in the *Nan* mice.

Understanding the role of KLF1 during erythroid maturation and the enucleation process has clinical significance for the production of red blood cells *in vitro* for transfusion purposes. In recent years, many efforts have been made to produce erythrocytes *in vitro* starting from hematopoietic stem cells, embryonic stem cells or induced pluripotent stem cells (42-44). Efficient enucleation is one of several challenges that have to be overcome in order to produce sufficient numbers of fully functional erythrocytes *in vitro*. More in depth knowledge of this process might guide the development of improved strategies to achieve this goal.

## Acknowledgements

TvD and HvdW were supported by the Netherlands Genomics Initiative (NGI Zenith 93511036); IC, TvD, FG and SP by the Landsteiner Foundation for Blood Transfusion Research (LSBR 1040); SH, MvL and SP by the Netherlands Organization for Health Research and Development (ZonMw TOP 40-00812-98-12128); IC, TvD, FG and SP by the EU fp7 Specific Cooperation Research Project THALAMOSS (306201). The funders had no role in study design, data collection and analysis, decision to publish, or preparation of the manuscript.

**Author contributions**
I.C., T.v.D,. H.J.G.v.d.W, R.S., M.v.L., F.G. and S.P. designed the experiments. I.C., N.G, R.S., S.H., A.M. and T.v.D. performed the experiments. H.J.G.v.d.W analyzed the 4C-seq and RNA-seq data and wrote the bioinformatics section. Z.O. and W.F.J.v.IJ. performed next generation sequencing experiments. The paper was written by I.C., H.J.G.v.d.W, M.v.L., F.G., T.v.D. and S.P. The authors have no conflicts of interest to declare.

